# Multivalent binding proteins can drive collapse and reswelling of chromatin in confinement

**DOI:** 10.1101/2022.01.21.477199

**Authors:** Sougata Guha, Mithun K. Mitra

## Abstract

Collapsed conformations of chromatin have been long suspected of being mediated by interactions with multivalent binding proteins, such as CTCF, which can bring together distant sections of the chromatin fiber. In this study, we use Langevin dynamics simulation of coarse grained chromatin polymer to show that the role of binding proteins can be more nuanced than previously suspected. In particular, for chromatin polymer in confinement, entropic forces can drive reswelling of collapsed chromatin with increasing binder concentrations. The reswelling transition happens at physiologically relevant binder concentrations and the extent of reswelling is mediated both by the concentration of binding proteins as well as the strength of confinement. We also study the kinetics of collapse and reswelling and show that both processes occur in similar timescales. We characterise this reswelling of chromatin in biologically relevant regimes and discuss implications for the spatial organisation of the genome.

## Introduction

The physical principles behind the three-dimensional organisation of the genome remains an interesting question in cell biology. The packaging of the chromatin polymer inside the confined space of a nucleus requires organisation and compaction at multiple hierarchical scales, while respecting topological constraints and being mediated both by the function of individual genes, and the genome as a whole. The organisation of chromatin inside the eukaryotic nucleus is a complex process, mediated by interactions of the DNA with multiple proteins. The organisation is dynamic[1, 2, 3], and changes in response to the environmental signals[4, 5], epigenetic state markers[6, 7], and cell cycle stages[8, 9], among others.

One of the structural motifs of this structural organisation is the formation of long-range contacts, mediated by proteins such as the CCCTC-binding factor (CTCF) [10, 11, 12, 13] and other proteins of the Structural Maintenance of Chromosomes (SMC) family, such as cohesin or condensin[14, 15, 16, 17]. In addition, many transcription factors are known to contain multiple chromatin binding domains, and can thus aggregate multiple nucleosomes [18]. The importance of long-range contacts on the 3D organisation of chromatin has been recognised for a long time. Loops, mediated by different binding proteins, brings together distant segments of chromatin, and plays an important role in the structural organisation of the genome [19, 20, 21], as well as in regulating gene expression [22]. Modifications in the structural organisation of chromatin has also been associated with various diseases [23, 5].

Experiments such as Hi-C and its variants have yielded a wealth of genome-wide data about long-range contacts [24, 25, 26, 27], and various polymer models have been proposed to integrate these experimental observations into a fundamental understanding of the physical principles of genome assembly[28, 29, 30, 31]. Simple self-avoiding polymer models have been used to reproduce different aspects of chromosome organisation, such as the emergence of chromosome domains and compartments [32, 33], Topologically Associated Domains (TADs) [34, 35], interactions of chromatin with the nuclear lamina [36, 37], and to understand the role of looping proteins in the formation and maintenance of chromatin loops [38].

Chromatin folding in presence of diffusible factors have been studied extensively. A widely used class of models for describing chromatin organisation inside nucleus is known as the “*Strings and Binders Switch*” (SBS) model [32, 39]. In this model the chromatin is assumed to be a self-avoiding polymer which consists of binding sites for diffusible factors (binders). These binders can generate different stable configurations of chromatin polymer through attachment. The chromatin polymer is found to undergo a coil-globule like transition from an open state to folded state as the binder concentration is increased [32, 40, 41]. The contact probability exponents in these studies are consistent with experimental observations [42]. On the other hand, stiffness of chromatin polymer and switching of binders between ‘on’ state and ‘off’ state play an important role in determining the structure of the clusters of chromatin formed by binders, and their kinetics [43, 44, 45]. Bridging by different multivalent transcription factors has shown separate clusters to form by different transcription factors and hence generates TADs [46]. Implicit bridging through direct interactions between monomers depending on their epigenetic states has also been studied [47, 48]. Other than forming different chromatin structure like loops, rochettes, TADs etc., distribution of epigenetic patterns on chromatin leads to coexistence or compartmentalization of different polymer domains (polymer-polymer phase separation).

A striking feature of chromatin organisation is the presence of structural and functional compartments within the nucleus [22, 49, 50]. At the largest scales, the nuclear volume confines the space available to chromosomes. However, confinement occurs at multiple smaller scales as well. Each chromosome occupies its own distinct space, known as chromosome territories [42, 51]. Within a chromosome, as revealed by Hi-C experiments, TADs represent megabasepair long regions enriched in internal contacts which play a key role in gene regulation [33, 52, 53]. The boundaries of these domains have been shown to be associated with CTCF protein [13]. Confinement of chromosomes has been recognised to be a potential regulator of both chromosome structure [36, 54] and dynamics [55]. In recent years, much research has focused on dynamic phase separated condensates in the nucleus, such as transcription factories associated with RNA polymerase II [56], and heterochromatin foci [57].

While the role of DNA-binding protein complexes in driving compaction of the chromosome has been studied in detail, these studies neglect the role of confinement. Multivalent counterions are known to drive non-monotonic behaviour of polymer sizes in polyelectrolytes [58, 59, 60], and multivalent DNA-binding proteins can play a similar role within the nucleus at higher concentrations. Compartmental organisation of nuclear material enable large fluctuations in concentration of DNA-binding proteins through local accumulation. Chromatin conformations can depend sensitively on such confinement induced fluctuations. In this work, we systematically investigate the coupled effects of DNA binder proteins and confinement of chromatin conformations, and show that chromatin-binder interactions can lead to rich and non-trivial behaviour of interphase chromosomes.

## Model and Methods

In this paper, we implement SBS model by assuming the chromatin as a self-avoiding flexible polymer consisting of 256 monomers. The bead (monomer) size is assumed to be greater than the Kuhn length of a persistent chain. The interaction between the polymer beads can simply be expressed as the sum of one repulsive and one attractive interaction [61]. The attractive repulsion is modelled via finitely extensible nonlinear elastic (FENE) bonds which ensures the finite extension of monomer bonds whereas the repulsive interaction, defined via Weeks-Chandler-Anderson (WCA) potential [62], incorporate the mutual volume exclusion. The FENE potential is given by,

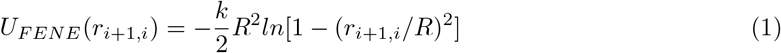

where *k* is the bond strength, *R* is the maximum possible distance between two consecutive monomers and *r_i,j_* = |*r_i_* – *r_j_*| is the relative distance between the i-th and j-th monomers.

The WCA interaction between the beads is given by,

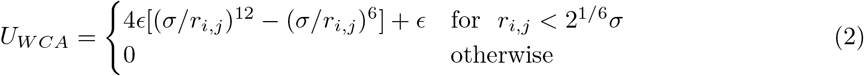

The FENE potential parameters are set to *k* = 30*ϵ*/*σ*^2^ and *R* = 1.6*σ* and the WCA interaction strength is taken to be *ϵ* = *k_B_T*.

The binder and monomer interaction is maintained by truncated and shifted Lennard-Jones potential and is given by,

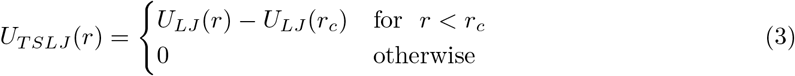

where *U_LJ_*(*r*) = 4*ϵ_m_*[(*σ*/*r*)^12^ – (*σ*/*r*)^6^] with *ϵ_m_* = 3.5*ϵ* and *r_c_* = 1.5*σ*. Motivated by experiments on binding valency of CTCF, and in line with previous modeling, we choose a binding valency of six for the binder proteins [63, 32]. A binder-monomer bond is considered to have formed instantaneously if they are separated by a distance less than *r_c_* and the binder is attached to less than 6 monomers. The binders interact among themselves via same WCA interaction given by Eq.2. The confinement is modelled as a soft wall defined again by the same WCA potential. The separation between the wall and the particle (monomer bead or binder) is measured from the closest point of the wall to the particle.

In our simulation, the energy and length is scaled by *ϵ* and *σ* respectively. We choose confinement radii to conform with the biologically relevant chromatin density inside nucleus. The human nuclear size varies in between 6-11 *μm* in diameter [64]. The human genome is known to consist 6 × 10^6^ kilobasepairs (kbp) [64] while 30 nm fibre bead is assumed to have 3 kbp [65]. Therefore we vary the confinement radii to maintain chromatin volume fraction (*ϕ*) in between 0.01 to 0.15 which is consistent with the known volume fraction of the chromatin. On the other hand CTCF abundance in human cells is found to be of the order of 10^5^ per cell [66] and hence suggest that CTCF molecules have a number density (*c*) in the range ~ 140 – 900 *μm*^-3^ inside nucleus. We assume *σ* ~ 70 nm in our simulation to conform with the experimentally observed CTCF number densities in cell. The timescale is scaled with the usual LJ time 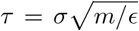 [67]. The mass of the particles are chosen to be unit, *m* = 1. The temperature of the system is maintained at *T* = *ϵ/k_B_* and the friction constant is taken to be *γ* = *k_B_T/τ*. It is easy to gain an intuition of the physical timescale (*τ*) in our simulation. The friction coefficient for spherical beads of diameter *σ* is given by *γ* = 3*πησ* where *η* is the nucleoplasm viscosity which is typically of the range of 10-100 cP [45]. Hence for typical value of *T* = 300 K, the timescale ranges in between *τ* = 3*πησ*^3^/*ϵ* ≃ 6 – 60 ms. We performed Langevin dynamics simulation using Velocity-Verlet algorithm [68] with time step *δt* = 0.002*τ*. In our simulation we observed all features of chromatin architecture to appear with CTCF (binder) numbers much less than the experimentally obtained number and within the CTCF density range consistent with experimental data [66] as well as previously reported theoretical studies [32, 40, 41, 44]. Thus even with larger polymer or confinement radius we expect all the features to be present within the biological range of CTCF number inside cell.

## Results

### A reswelling transition of chromatin

We first characterise the equilibrium properties of the chromatin polymer for different compaction levels and with differing binder numbers. For the largest radius of confinement (*R* = 15, *ϕ* = 0.01), as shown in Fig. 2a, in the absence of binders the polymer adopts a semi-open conformation (*R_g_* ≃ 8.1, 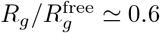, where 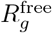 denotes the radius of gyration for the unconfined polymer). As binders are introduced, the polymer collapses due to binder-mediated long-range interactions between distant polymer segments. For *R* = 15, the minimum radius of gyration, 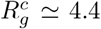 is achieved at a critical binder number of 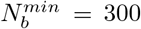 with number density *c^min^* ≃ 0.02. Contrary to existing models of binder mediated collapse however, on further increasing the number of binders, the polymer does not stay in the maximally collapsed conformation, but starts to reswell with increasing binder numbers. The extent of reswelling depends on the number of binders, but also on the confinement radius. For *R* = 15, where the polymer volume fraction is low, the collapse occurs much faster, followed by a slow reswelling. This generic nature of a collapse from a semi-open state to a compact conformation, followed by a reswelling on further increasing binder concentrations is observed at all values of the confinement (Fig. 2b for *R* = 9, and Fig. 2c for *R* = 6.5). The critical concentration required for the collapse of the polymer increases with increasing confinement, since a certain minimum number of binders are required to form enough long-range contacts for effective collapse, which translates to higher concentration for highly confined systems. Note that with increasing confinement, the polymer is increasingly compact even in the absence of binders, and hence the relative change in *R_g_* from the zero-binder case to the maximally compact conformation decreases with stronger confinement. Conversely, the rate of reswelling with increasing binder numbers (or concentration) is much higher for stronger confinement, with radius of gyration reverting to the zero-binder value for *R* = 6.5. In order to take into account this differential degrees of collapse for different values of the confinement radius, we define a compaction parameter,

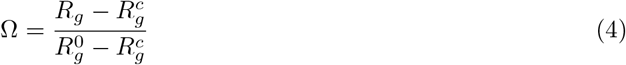

where 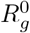 and 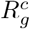 are the radius of gyration of the polymer in free and fully compacted states respectively. Ω = 1 then denotes a free state (i.e no binders present) and Ω = 0 corresponds to the fully compacted state (i.e minimum *R_g_*). The variation of the compaction parameter as a function of the concentration for all three values of the confinement radius is then shown in Fig. 2d. As discussed above, for weakly confined systems, the collapse transition occurs at lower concentrations, however, the reswelling as a function of binder concentration is sharper for strongly confined systems. Note that the concentrations at which reswelling is appreciable are within biologically relevant regimes, and hence this counter-intuitive reswelling is expected to play a role within the nucleus.

**Figure 1:**
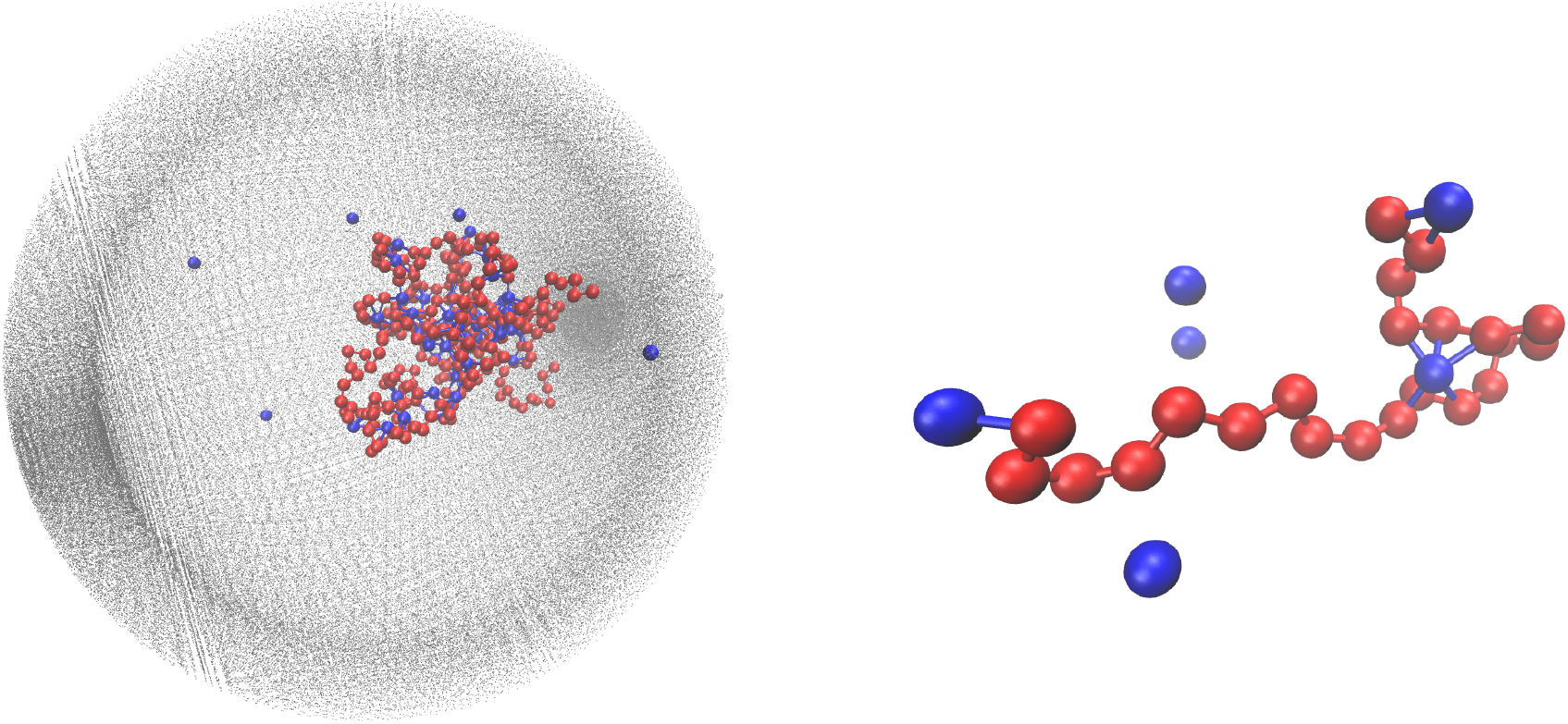
Full scale schematic (left) of a polymer in confinement in the presence of binding protein. A blown-up illustration of the chromatin-binder interactions is shown on the right. Red beads depict the monomers and the blue beads represent the binders. The monomer-monomer bonds are shown by red solid lines while monomer-binder bonds are shown by blue solid lines. The 3D confinement is portrayed by grey dots.

**Figure 2:**
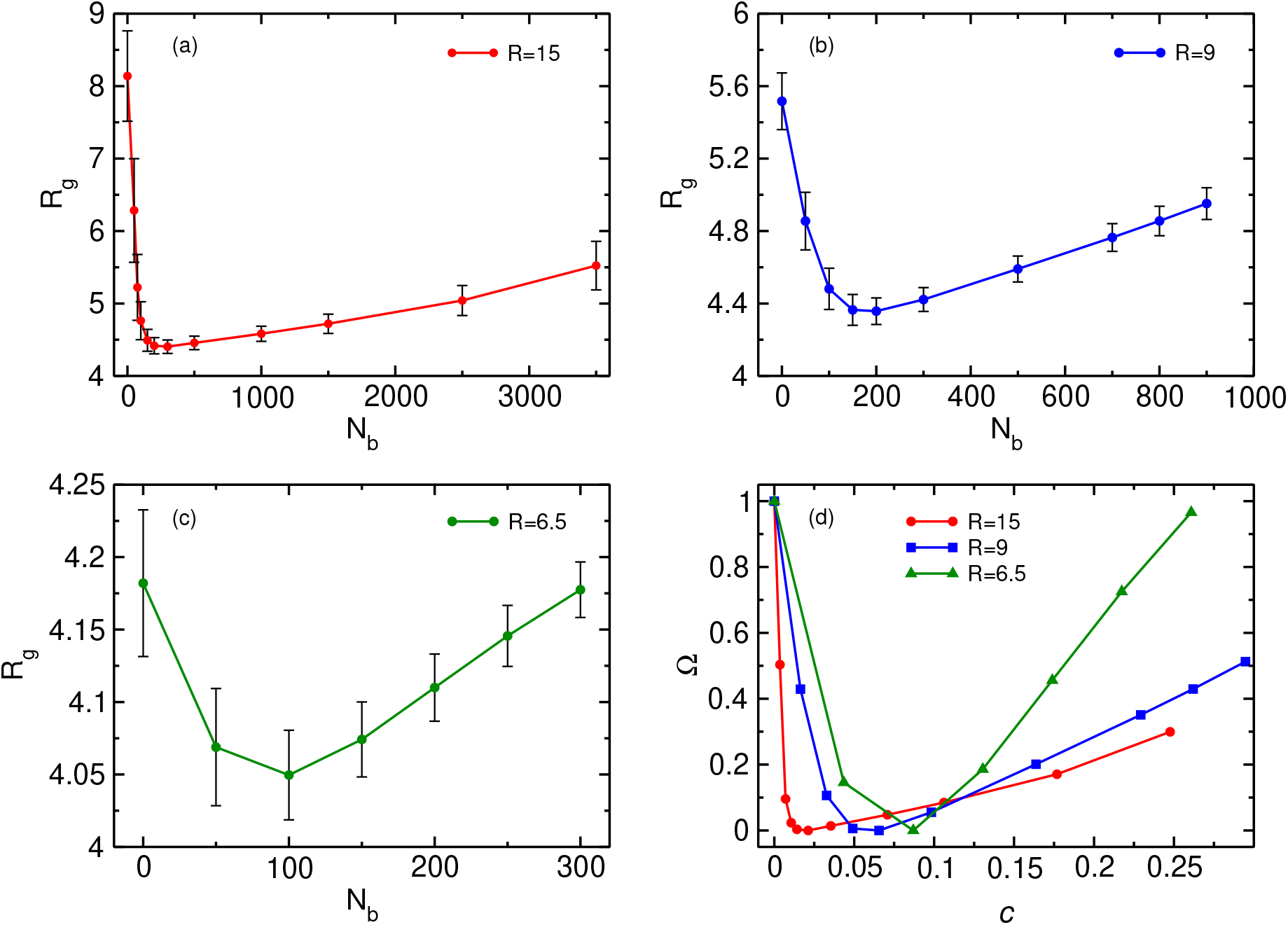
The plot depicts (a-c) the variation of radius of gyration (*R_g_*) with binder number and (d) the variation of compaction parameter Ω with binder number density for 3 different confinement radii *R*.

### Binder mediated reswelling is driven by competition between energy and entropy

In order to understand the role of binders in the collapse and reswelling of the polymer, we plot the probability *p_m_*(*j*) of having *j* monomers bound to a single binder protein (Fig 3a), and the probability *P_b_*(*j*) of having *j* binders bound to a single monomer (Fig. 3b), for different binder numbers. The plots are shown for the intermediate confinement case, *R* = 9, but the general trend holds true for any strength of confinement. When the number of binders is very low (*N_b_* = 50), the polymer is in a semi-open conformation (Ω = 0.5), and most binders have the maximum possible number of monomers bound to them, resulting in a distribution of *p_m_* peaked at 6 monomers. However, because of the low number of binders, most monomers have none or one binder bound to it (Fig. 3b), resulting in very few long-range contacts, as expected for this semi-open conformation. In this conformation state therefore, most binders are surrounded by a cloud of monomers, as shown in Fig. 3c. If we now consider the case of the critical binder number for which we observe the most compact conformation, 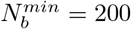, the distribution of the number of monomers bound to a single binder is qualitatively similar to the *N_b_* = 50 case, with most binders saturating their maximum valency of 6 monomers. However, for this compact conformation, the *p_b_* distribution undergoes a significant change from the low binder regime, with a distribution peaked at around 4 binders, and very few monomers remain which have at most one binder bound to them. At this critical binder number therefore, the polymer-binder system achieves an optimum for the compact conformation. On increasing binder numbers past this critical value, there is now an excess of binders, which results in a drastic change in the *p_m_* distribution, with very few binders now achieving the maximum valency. Conversely, the distribution of *p_b_* shifts even further to the right, resulting in each monomer acquiring a cloud of binders surrounding it, as shown in Fig. 3d. Since each monomer is now surrounded by a binder cloud, binders no longer require long-range contacts to minimise their energy, and hence the polymer reswells as it becomes more entropically favorable. Further, the binder cloud surrounding a monomer increases the effective size of a single monomer, and hence excluded volume forces also contribute to the reswelling.

**Figure 3:**
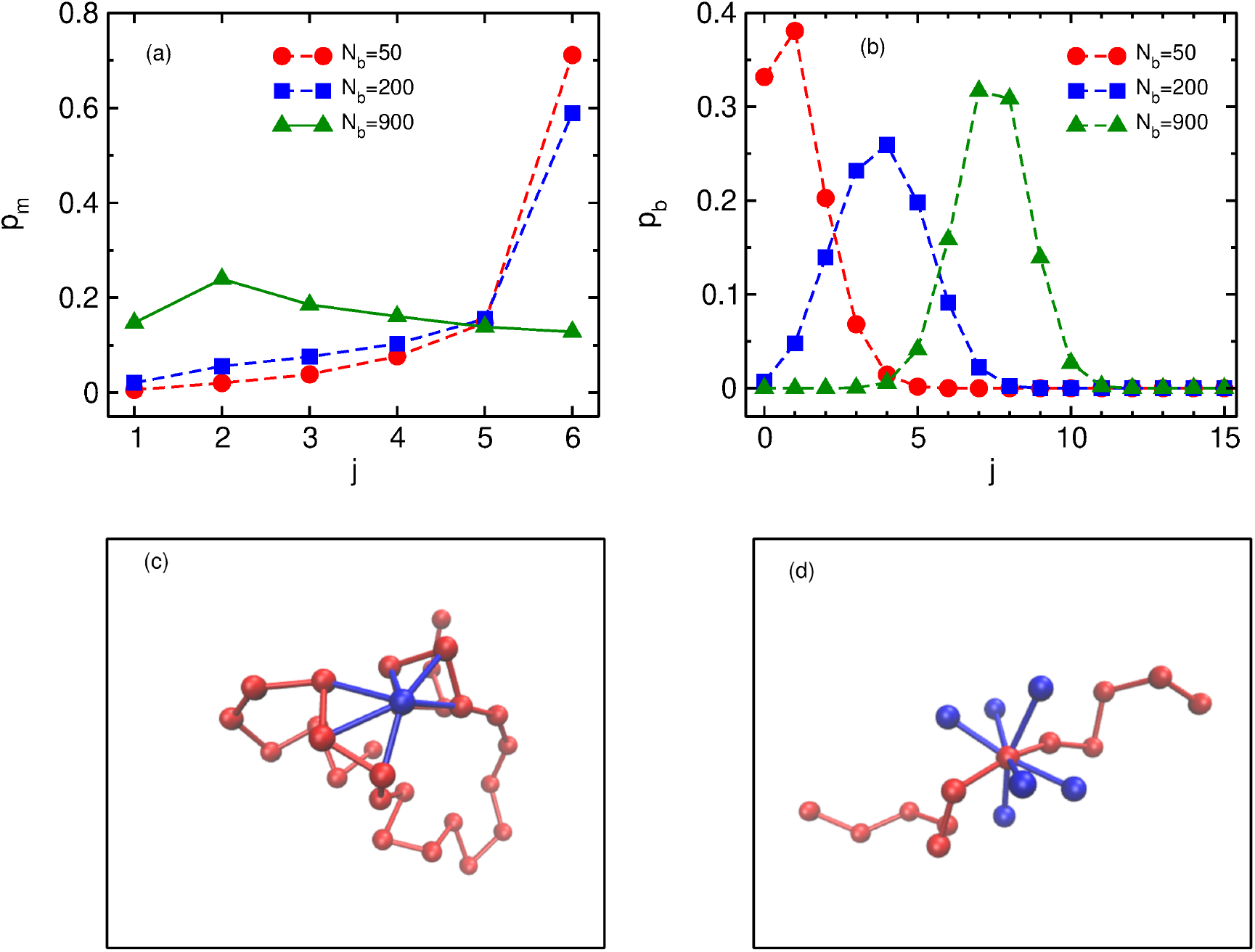
The plot depicts the probability distributions of (a) monomers bound to a single binder (*p_m_*) and (b) binders bound to a single monomer (*p_b_*) for 3 different binder numbers corresponding to semi-open, compact and reswelled chromatin for *R* = 9. Panels (c-d) show representative figures of monomer cloud around binder and binder cloud around monomer respectively.

We now turn to the contact probability distributions in order to understand the coupled effects of binder concentration and confinement for the (semi-)open, compact and reswollen conformations. Generically, the contact probability scales with genomic distance as, *P_c_*(*s*) ~ 1/*s^α^*, where the exponent *α* in this case depends on both the binder concentration as well as confinement. We first consider the case of weak confinement, R = 15, shown in Fig. 4a. In the absence of binders, the contact probability exponent goes as α ~ 2.0, indicating the polymer conformation is similar to the open random SAW polymer. Close to the transition point for the collapsed conformation, 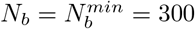, the contact probability exponent is *α* ~ 0.75, which is in the biological range observed in experiments. On increasing the binder concentrations (*N_b_* = 3500), the polymer reswells, which is reflected in a corresponding increase in the exponent, α ~ 0.9, indicating a decreased probability of contact between distant monomer segments. Note that for this case of weak confinement, biological constraints on the binder concentrations preclude a full reswelling of the polymer. In order to understand the equilibrium properties of the conformations of this partially reswollen conformation, we then compare it to the conformation at a pre-collapse binder concentration, 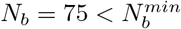, such that the radius of gyration of these two conformations are roughly similar (*R_g_*(*N_b_* = 3500) ≈ *R_g_*(*N_b_* = 75) ≃ 5.3). Firstly we note that for small genomic separations (*s/σ* < 10), the contact probabilities for the reswollen polymer are lower than the partially collapsed polymer, which can be understood by the presence of the binder cloud surrounding monomers in the reswollen case, as discussed above. At larger separations however, the contact probability exponent in both these cases are roughly similar, indicating that reswollen conformations are statistically similar to the open counterparts. For the case of intermediate confinement, *R* = 9, again in the absence of binders, the contact probability behaves similar to a semi-open conformation (*α* ~ 2.0 for *N_b_* = 0). At the collapsed conformation, the contact probability exponent becomes much smaller, *α* ~ 0.9, reflecting the increased long-range contacts in this collapsed conformation. As the polymer reswells (*N*_b_ = 800), the contact probability itself becomes lower due to the presence of a binder cloud surrounding each monomer, although the exponent *α* in this case is roughly similar to the compact conformation. Similar features are observed for *R* = 6.5, although the determination of the exponent in this case is complicated due to the strong confinement induced collapse.

**Figure 4:**
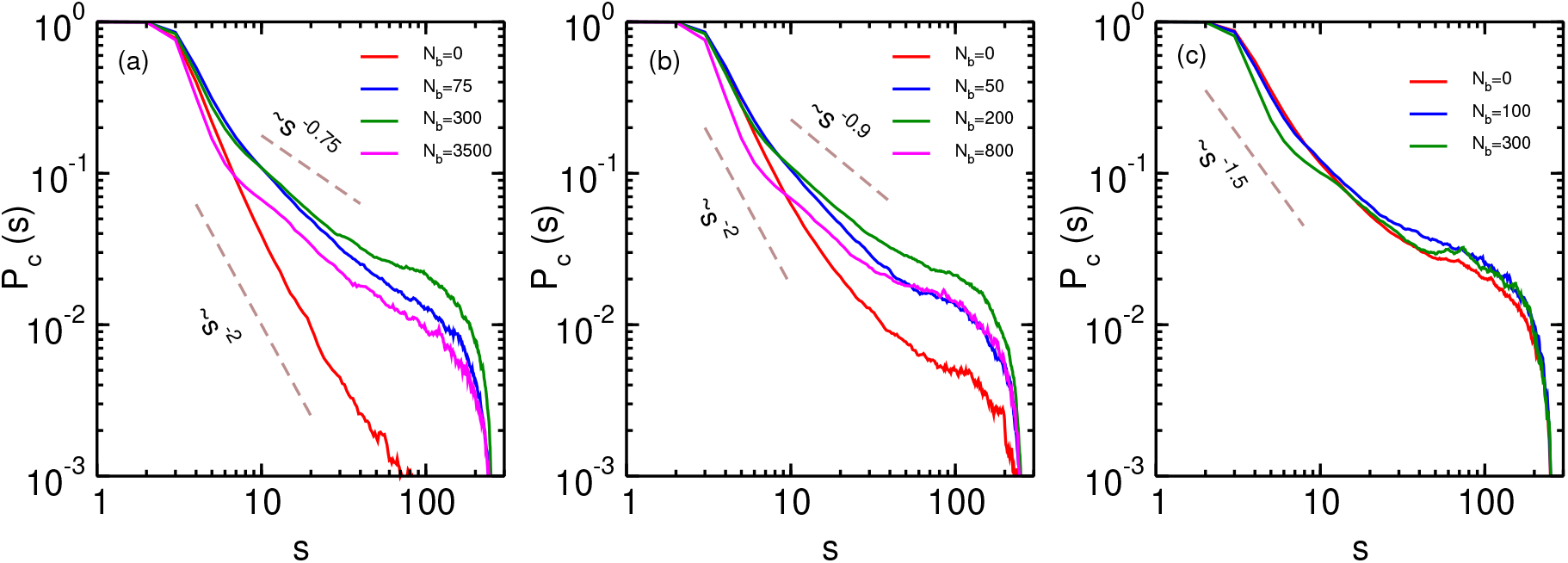
Contact probability of the polymer with varying binder numbers for 3 different confinement radii R. Panel (a) corresponds to R = 15, panel (b) corresponds to R =9 and panel (c) corresponds to R = 6.5. Dashed lines show guides to the eye for different values of the contact probability exponent.

### The kinetics of collapse and reswelling

We now turn to the kinetics of the collapse and the reswelling of the chromatin polymer in order to investigate whether both of these processes can occur within biological timescales. We first look at the distributions of compaction times. At a given value of the confinement radius R, we allow the chromatin polymer to equilibrate in the absence of any binders (*N_b_* = 0). We then instantaneously increase the number of binders to the critical binder number corresponding to maximum compaction 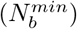, and monitor the time evolution of the radius of gyration of the polymer (*R_g_*(*t*)). The compaction time *τ_c_* is defined as the minimum time at which *R_g_*(*t*) reaches the equilibrium value 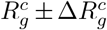, *and* the time-averaged < *R_g_* >*_t_* remains within this equilibrium range for the next 10^5^(200*τ*) steps. The distributions (*p_c_*) of this *τ_c_* for three different compaction radii are shown in Fig. 5a-c. The distributions of the compaction times are unimodal, and can be approximated by log-normal distributions. The mean compaction times increase with increasing radius of confinement, and are given by

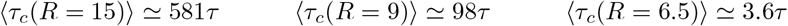

**Figure 5:**
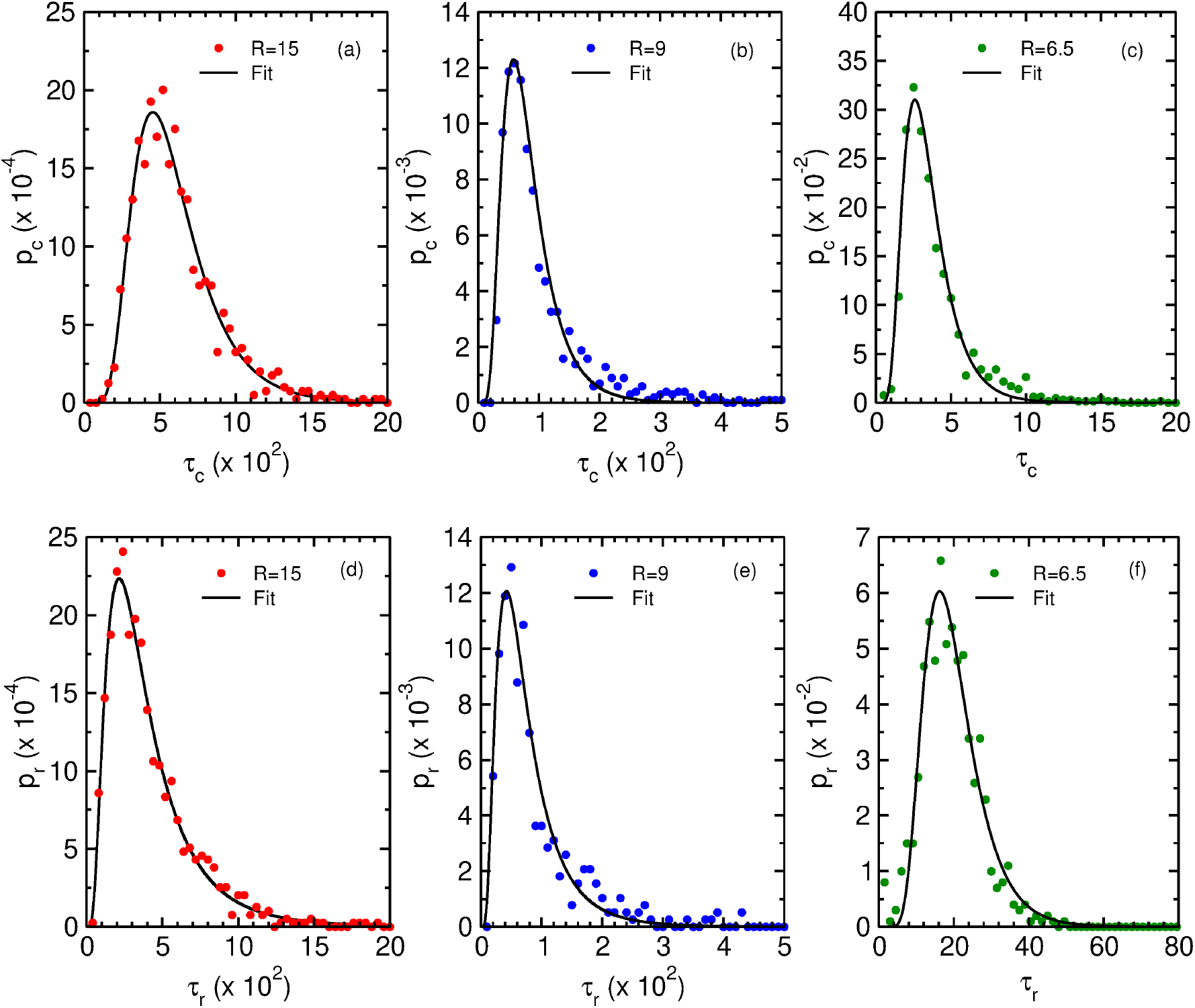
The plot depicts (a-c) the compaction time distribution *p_c_* as a function of *τ_c_* and (d-f) reswelling time distribution *p_r_* as a function of *τ_r_* for different confinement radii *R*. The dots represent the simulation results and the black line represents a log-normal fit of the simulation data. The *σ* and *μ* values of the fit are (a) *σ* ~ 0.43, *μ* ~ 6.3, (b) *σ* ~ 0.49, *μ* ~ 4.3, (c) *σ* ~ 0.45, *μ* ~ 1.15, (d) *σ* ~ 0.66, *μ* ~ 5.81, (e) *σ* ~ 0.64, *μ* ~ 4.15, (f) *σ* ~ 0.38, *μ* ~ 2.93. The reswelling time distributions are obtained with binder density *c* = 0.2 for all radii.

For larger values of the confinement radius, the change in the radius of gyration between the open and compact states is larger and hence leads to larger compaction times. Note that with a typical timescale of *τ* ~ 10*ms*, these collapse timescales range from tens of milliseconds to a few seconds, well within biologically relevant ranges.

We next ask whether binder mediated reswelling also occurs within physiological timescales. To check this, we initialise our system in the compact conformation, by equilibrating the chromatin polymer with 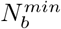 binders. We then instantaneously increase the binder concentration to *c* = 0.2 for all values of the confinement radii *R*. This induces a reswelling of chromatin, and the reswelling time *τ_r_* is defined similarly to *τ_c_* by noting the minimum time taken to reach the equilibrium *R_g_* and then fluctuate around this value for 200τ steps. The distributions of the compaction times are again unimodal, and well-approximated by log-normal distributions (Fig. 5d-f). Note that since the concentration of binders is kept uniform, the extent of reswelling is different for different *R* (Ω_15_ = 0.2, Ω_9_ = 0.3, Ω_6.5_ = 0.7). For the case of maximal confinement, when c = 0.2 approaches the maximum possible reswelling (Ω ~ 1), we note that the mean reswelling time ⟨*τ_r_*⟩ ≃ 5⟨*τ_c_*⟩. Irrespective of the extent of reswelling, we note that the mean reswelling times are comparable to the collapse timescales, suggesting that modulating local binder concentrations can lead to collapse and expansion of polymers within biological timescales.

### The effect of confinement radius on collapse and reswelling

Local binder concentrations in cells can be sensitively controlled by changing the effective confinement radius. In this section, we study the effect of varying confinement radius on the collapse and reswelling timescales. We first characterise how mean collapse times *τ_c_* change with changing confinement. For a given value of the confinement radius (*R*), the extent of collapse can be identified as the difference between the binder-free radius of gyration 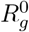 and the final compaction value 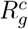 both of which depends explicitly on *R*. While 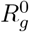 depends strongly on *R*, the variation of 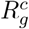 with confinement radius is comparatively smaller indicating a weaker dependence of 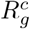 on *R*. Therefore we use 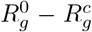 as a proxy for variation of confinement radius to determine how compaction time depends of confinement. We find that the mean collapse times scales with the extent of compaction as 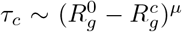 with *μ* = 1.45 ± 0.06 (Fig. 6a). Previous studies, in the absence of confinement, have reported compaction time scaling with polymer size with a similar exponent, *τ_c_* ~ *N*^1.5^ [69]. Confinement has a non-trivial effect on the scaling of polymer *R_g_* with polymer size [55], and hence the extent of confinement 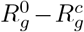 provides a more robust characterization of the compaction timescales.

**Figure 6:**
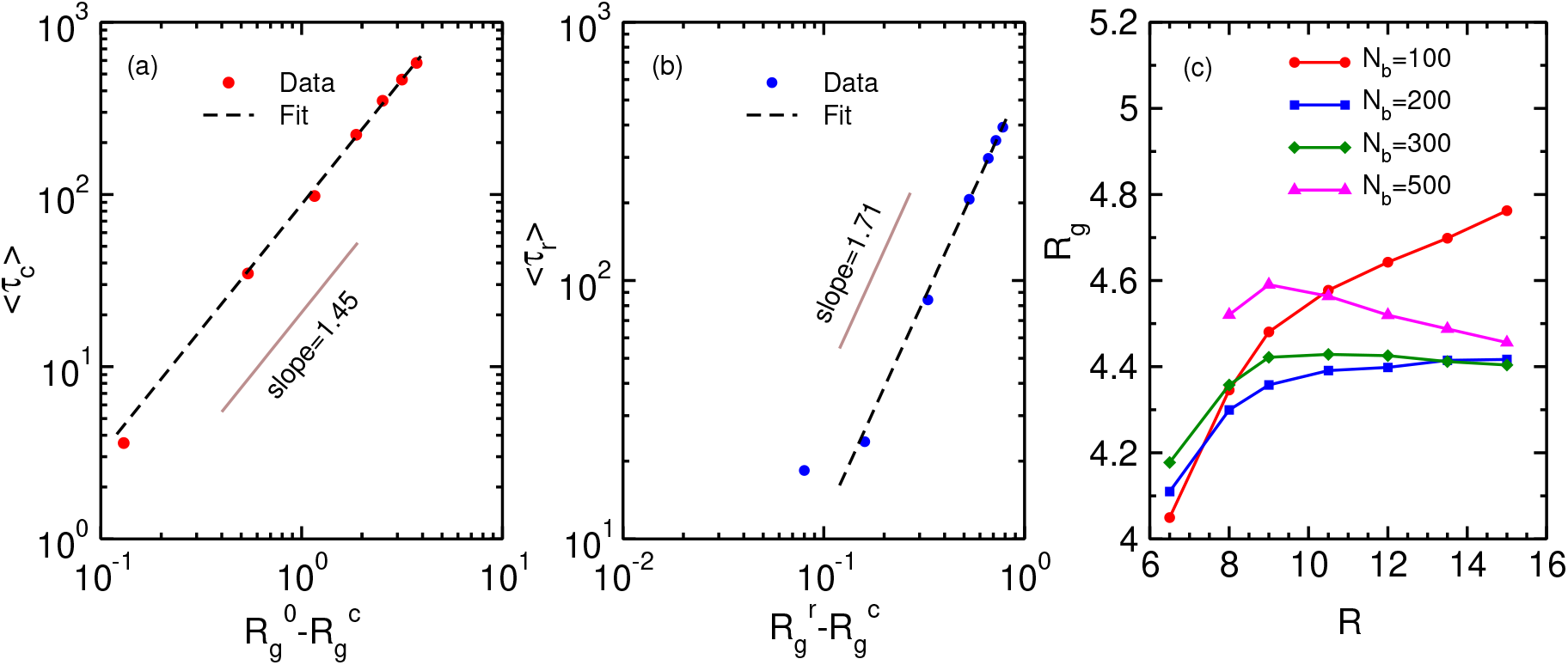
The plot depicts the scaling of (a) mean compaction times (⟨*τ_c_*⟩) with the amount of compaction in polymer size, and (b) mean reswelling time (⟨*τ_r_*⟩) with the extent of reswelling in polymer sizes. The mean compaction time (*τ_c_*) scales with the difference between zero-binder radius of gyration 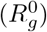 and compacted radius of gyration 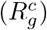 with exponent *μ* ≃ 1.45 whereas the mean reswelling time (*τ_r_*) scales with difference between reswelled radius of gyration 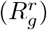 and compact radius of gyration 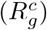 with exponent *v* ≃ 1.71. The different values of 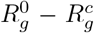 (panel a) and 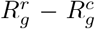 (panel b) correspond to different values of the confinement radius *R*. Panel (c) shows the variation of radius of gyration (*R_g_*) with change in confinement radius (*R*) for 4 different N_b_ values.

Similarly, we also characterize the scaling of the mean reswelling time with changing confinement radius. For a given confinement radius, we characterize the extent of reswelling by the difference between the reswelled radius of gyration 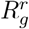 and the compact radius of gyration 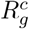. Therefore, similar to compaction times, in this case we use 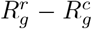 as the proxy for variation of confinement. The reswelling time scales with the extent of reswelling as 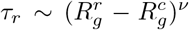 with *v* = 1.71 ± 0.07 (Fig. 6b). The reswelling exponent is thus slightly higher than the compaction exponent (*vμ*), and for all biologically relevant values of 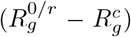, and equivalently of the confinement radius, the compaction and reswelling times are expected to be similar in magnitude, and hence both processes are expected to be physiologically relevant.

We now turn to the control of polymer size (*R_g_*) by changing the confinement radius, and hence the local binder concentration. This is shown in Fig. 6c for different values of the binder number *N_b_*. For low number of binders, when the chromatin polymer is in the semi-open state, it is expected that the polymer is distributed over the whole available volume, and hence *R_g_* is expected to increase with increasing volume (and equivalently *R*). Evidently we found this to be true in presence of less number of binders (*N_b_* = 100). As the binder number is increased (*N_b_* ~ 200 – 300), *R_g_* tends to saturate with increasing confinement radius. This is due to the fact that the binder number is close to the critical binder number 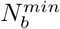 for larger values of the confinement radii and the polymer collapse into a compact state in these cases. Further increase in binder numbers results in a re-entrant like transition (*N_b_* = 500). For strong confinement, *R_g_* initially increases with increasing *R*. However, beyond a certain confinement radius, *R_g_* starts to reduce with increasing *R*. This can be explained in terms of reswelling of the chromatin. For strong confinement *N_b_* = 500 is significantly larger than 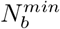 corresponding to those confinement radii and thus the polymer is in reswollen state. On the other hand for large confinement radii *N_b_* = 500 is close to 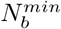 for weak confinements and therefore the polymer is still in a moderately compact state resulting in lower *R_g_* values. This non-trivial re-entrant behavior shows how the cell can control chromatin sizes by modulating the extent of confinement, for the same number of binding proteins.

## Discussion

We show, that contrary to existing models of binder protein mediated collapse of chromatin conformations, chromatin polymer configurations depend sensitively on binder concentrations, and high concentrations of binders can in fact drive reswelling of chromatin. Our simulations show that while at low concentrations binders mediate collapse by forming loops between distant polymer segments which lowers the energy of these compact conformations, at larger concentrations there are enough binders that each monomer is surrounded by a binder cloud, and hence the polymer preferentially maximises entropy through semi-open swollen conformations. Additionally, we show that timescales of collapse are similar to timescales of reswelling, and hence both of these effects are expected to play an important role within cells.

The coil-globule transition of chromatin in unconfined space has been well studied in the literature [32, 40, 41, 42, 43, 45, 44]. While studies of confined polymers have shown novel features such as glassy dynamics upon strong confinement [55], effects of confinement on chromatin polymer in presence of chromatin binding proteins is has not been well studied. A recent study has postulated reswelling by bridging molecules in the absence of confinement, which however occurs only at very high volume fractions [70]. Our study suggests that the reswelling of the chromatin depends heavily on the confinement radius as well as presence of multivalent binding proteins, and the onset of reswelling appears at different binder density for different confinement radii even when the binder-chromatin interaction strength is unchanged.

Our simulations show that the chromatin sizes and compaction can be sensitively controlled by changing the confinement radius. This offers a tantalising probability for the control of formation and stability of phase-separated condensates in the nucleus [50]. Recent experiments have shown that CTCF is locally concentrated inside transcriptional condensates, and CTCF mediated looping is in fact an essential architectural prerequisite for the formation of such condensates [12]. This is consistent with our simulations. We find that multivalent binding proteins are locally enriched within the polymer phase (with an average concentration 10 – 100 times higher inside the collapsed polymer phase as compared to the surrounding fluid). Fluctuations in the size of the compartment can then push local concentrations above or below the reswelling threshold, with an accompanying change in the structure of the chromatin polymer.

Our course-grained model simulations, while it neglects sequence heterogeneity and other biological interactions, shows that in the presence of confinement, multivalent binding proteins can either stabilise or destabilise the collapsed polymer conformation through a non-trivial interplay between binder concentrations and confinement strengths.

## Author Contributions

M.K.M designed the research. S.G. developed and carried out all simulations. S.G. and M.K.M. analysed all data, interpreted results, and wrote the article.

## Acknowledgments

SG and MKM acknowledge IIT Bombay for research funds and support. MKM acknowldged IIT Bombay Grant No. 14IRCCSG009 for financial support.

